# Alpha power indexes working memory load for durations

**DOI:** 10.1101/2025.05.12.653390

**Authors:** Sophie K. Herbst, Izem Mangione, Charbel-Raphaël Segerie, Richard Höchenberger, Tadeusz Kononowicz, Alexandre Gramfort, Virginie van Wassenhove

## Abstract

Timing, that is estimating, comparing, or remembering how long events last, requires the temporary storage of durations. How durations are stored in working memory is unknown, despite the widely held view of memory systems’ central role in timing. Here, we investigated the neural signatures of a sequence of durations (n-item sequence) held in working memory. We recorded human participants using magnetoencephalography (MEG) while they performed an n-item delayed reproduction task, which required to encode a sequence of durations, maintain it, and then reproduce it. The number of items in a sequence (one or three) and the duration of the sequence were orthogonalized. Our results show that during working memory maintenance, the number of durations, not the duration of the sequence, affected recall precision and could be decoded from alpha and beta oscillatory activity. Parieto-occipital alpha power showed a direct link with the precision of temporal reproduction. Our results extend earlier behavioral findings suggesting that durations are itemized in working memory and that their number, not their duration, modulates recall precision (Herbst et al., 2025). Crucially, we establish that alpha power reflects a universal signature of working memory load and mediates recall precision, even for abstract information such as duration.

## Introduction

Humans and animals can readily discriminate the durations of sensory events as well as the silent intervals that may separate them. Sensory timing tasks typically quantify this ability by asking individuals to compare or to (re)produce time intervals (Grondin, 2010). In most sensory timing tasks, the encoded temporal information is stored until a decision is made: for instance, in the simplest two-alternative-forced choice design, two time intervals are presented in succession, and the first of the two is held in working memory until the second is presented and a comparison can be made (Coull et al., 2008; Harrington et al., 2010; Rao et al., 2001). The encoding of duration in such tasks and the decision stages have been well explored (Díaz et al., 2025; Ofir & Landau, 2022; Paton & Buonomano, 2018). To the contrary, how durations are maintained in working memory is unknown. Classical information-theoretic timing models postulated a working memory component (Buhusi & Meck, 2005; Gibbon, 1977; Gibbon et al., 1984; Treisman, 1963), but its mechanisms and neural dynamics remain largely speculative (Gu et al., 2015; Teki et al., 2017; van Wassenhove, 2016).

Recent behavioral work suggests that the storage of durations in working memory is comparable to that of any items (Herbst et al., 2025; Manohar & Husain, 2016; Teki & Griffiths, 2014), in that recall precision is indicative of working memory load (Bays, 2015; Bays et al., 2024; Ma et al., 2014). At the cortical level, the maintenance of encoded events in working memory has been linked to oscillatory dynamics (Miller et al., 2018), which provide a self-sustained neural code in the absence of sensory stimulation. Working memory function likely relies on different oscillatory regimes, ranging from theta to alpha, beta, and gamma (Axmacher et al., 2010; Jensen & Lisman, 2005; Kopell et al., 2011; Lisman & Idiart, 1995; Palva & Palva, 2011; Roux & Uhlhaas, 2014). A consistent finding is that alpha power (8 – 12 Hz) increases with working memory load (Jensen et al., 2002), but studies assessing working memory for duration do not clearly align with this seminal observation as discussed below (Y. G. Chen et al., 2015; Y. G. Chen & Huang, 2016).

Herein, we thus investigated whether duration storage follows the known principles of working memory. We focused on recall precision as a measure of working memory load, and neural oscillations as its neural signature. We recorded participants’ brain activity with magnetoencephalography (MEG) while they performed an n-items delayed reproduction task, in which they were asked to encode, maintain, and reproduce temporal sequences of varying length. Critically, we orthogonalized the number of items (one or three) in a sequence and the duration of the sequence. Using this task, we previously showed that the number of items, but not the duration of the sequence affected recall precision (Herbst et al., 2025). These results provided evidence that durations can be itemized and stored in working memory as mental abstract magnitudes (Bueti & Walsh, 2009; Gallistel, 2011; Gallistel & Gelman, 2000; Walsh, 2003). Accordingly, in the present study, we expected an increase in alpha power with working memory load depending on the number of durations in a sequence, but not on the duration of the sequences itself.

The modulation of alpha power is a well-established marker of working memory load, resulting from the number of items (Jensen et al., 2002; Klimesch et al., 1999; Scheeringa et al., 2009). Jensen et al. (2002) demonstrated a parametric alpha increase over posterior and parietal regions during a visual Sternberg task (2, 4, and 6 items, electroencephalography (EEG)), a finding that has since been replicated in visual (Ede et al., 2018; Kornblith et al., 2016; Tuladhar et al., 2007) and auditory contexts (Kaufman et al., 1992; Krause et al., 1996; Leiberg et al., 2006; Obleser et al., 2012; see also the review by Wilsch & Obleser, 2016). Working memory load is classically operationalized through the number of items that have to be maintained, but similar results have been obtained when modulating the complexity of the stimuli (X. Chen et al., 2023; Krause et al., 1995; Obleser et al., 2012)

The increase in alpha power is thought to reflect inhibition of task-irrelevant information during retention (Bonnefond & Jensen, 2012; see also Klimesch et al., 2007; Schneider et al., 2021), and can be modulated by temporal expectations about the retention interval (F.-W. Chen et al., 2023; Wilsch et al., 2014). Two studies have directly targeted the neural dynamics of working memory maintenance for duration: one study showed that alpha power decreased with longer visual item durations (Y. G. Chen et al., 2015), whereas the second study, using a visual n-back task, reported a decrease in posterior alpha power with increasing working memory load – together with beta power increase over temporal areas (Y. G. Chen & Huang, 2016). The divergent alpha power trends in timing studies suggest the possibility of distinct mechanisms for durations held in working memory. To investigate this further, we employed alpha power as an index of working memory load, aiming to determine which factor, the number of items (n-items) or the duration of the sequence contributes most to working memory load.

We show that the number of items, but not the duration of the sequence, can be decoded from both alpha and beta (15 – 25 Hz) band activities, with distinct cortical sources. Interestingly, the alpha-band power showed a direct link to the precision of temporal reproduction. Thus, our results suggest that durations are itemized in working memory like any other mental events.

## Results

### Behavioral Results

In an n-item-delayed-reproduction task, participants reproduced sequences of either one or three duration-items following a retention interval (see **Figure 1A**). The sequences varied orthogonally in the number of items (one or three), and the duration of the sequence (1.6 s / short, 2.4 s / medium, 3.6 s / long, **Figure 1B**). The behavioral results replicate our previous findings (Herbst et al., 2025). Specifically, the relative reproduction (relRP, defined as the ratio between each item’s reproduction and duration) was modulated by the number of items and the sequence duration (Table 1, Figure 1C, Supplementary Figure 1). Consistent with context effects in magnitude estimation (Petzschner et al., 2015), durations in sequences containing more items were reproduced as longer compared to durations that occurred alone. Furthermore, short sequences were overproduced, long sequences were underproduced, an effect typically interpreted as a regression to the mean (Lejeune & Wearden, 2009; Petzschner et al., 2015; Vierordt, 1868).

**Table 1.**
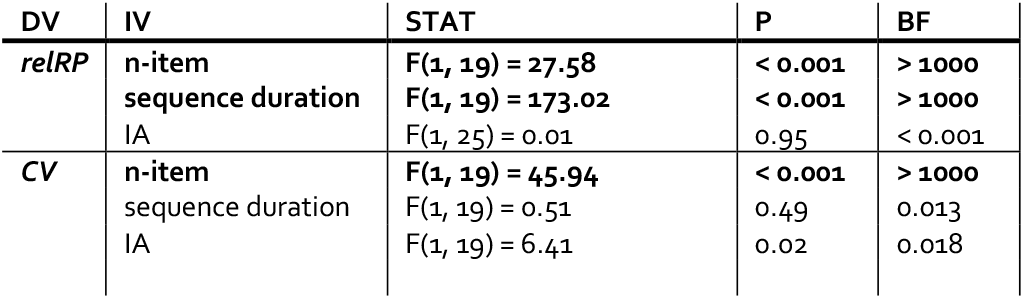
Effects of the number of items (n-item) and sequence duration on the relative reproduction (relRP) and precision (CV) of temporal reproductions. Bayes Factors (BF) reflect the Bayesian evidence for the significance of the respective predictor. Bold values indicate a significant effect. DV = dependent variable, IV = independent variable, STAT = statistical parameters, P = p-value, IA = interaction.

**Figure 1.**
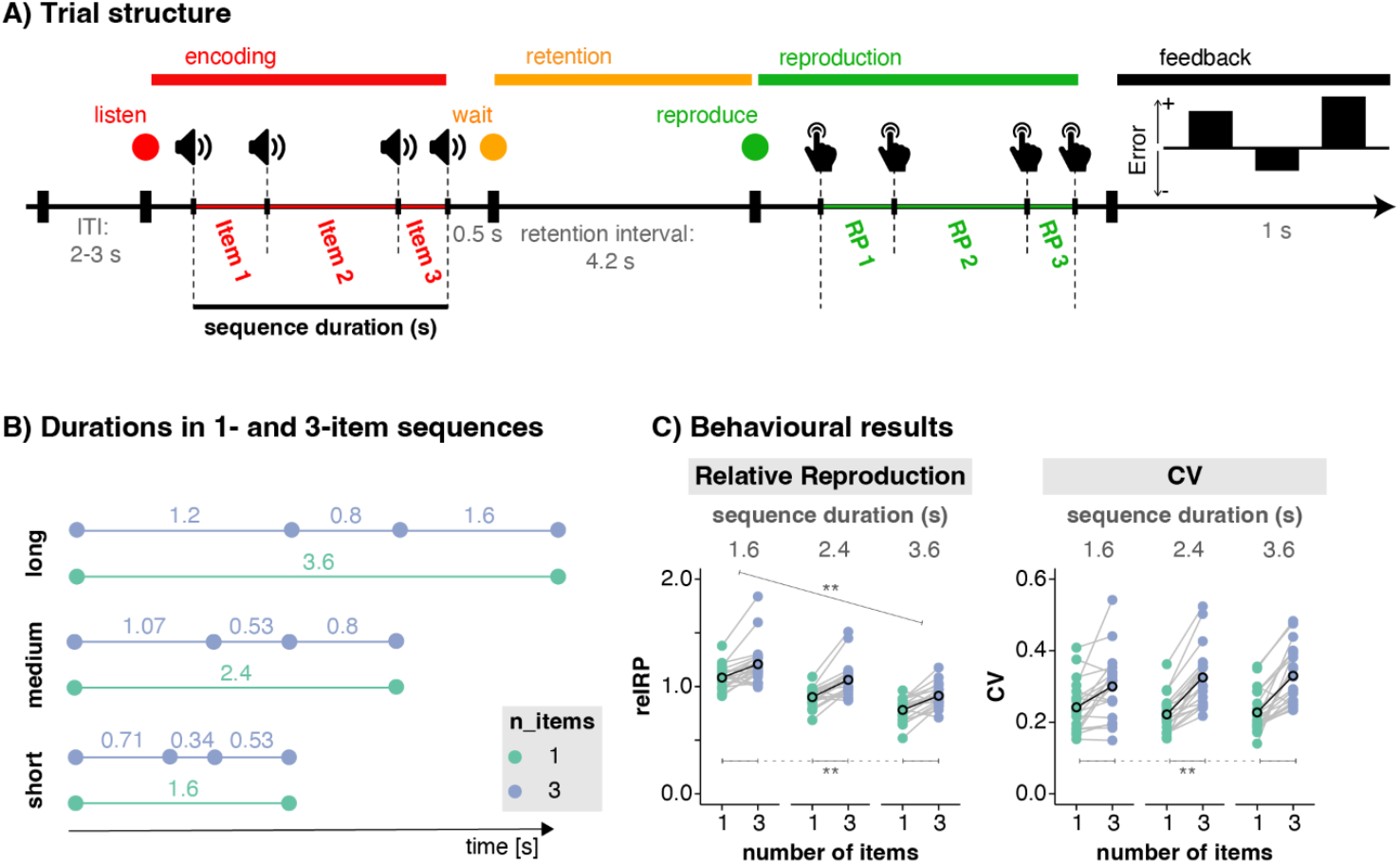
N-item delayed reproduction task and behavioural results. **A: Trial structure**. Each trial consisted of four phases: First, participants listened to a sequence of pure tones (1 kHz, 50 ms) demarcating empty time intervals of varying duration (encoding phase, red fixation dot). Following a retention interval (orange fixation dot), participants were asked to reproduce the full temporal sequence as precisely as possible (green fixation dot). In the example depicted here, the sequence was composed of three durations. Following their reproduction, participants received feedback about the relative temporal reproduction error for each duration in the sequence. ITI = inter-trial-interval, RP = reproduction. **B: Items and sequences**. Dots represent tones. Horizontal lines and the numbers above indicate the duration of the empty intervals in seconds. A sequence was composed of 1 or 3 items (blue and green, respectively). The durations of sequences were fixed to 1.6 s (short), 2.4 s (medium), or 3.6 s (long), irrespective of the number of items that composed them, such that the number of items and the duration of the full sequence were orthogonalized. **C: Effects of the number of items and the duration of sequences on relative reproduction (relRP) and inverse precision (coefficient of variation, CV)**. Left: relRP significantly increased with the number of items and decreased with the duration of the sequence (** = p < 0.01). Right: CVs were not affected by the duration of the sequence but significantly increased with the number of items, indicating a decrease in reproduction precision with higher working memory load.

More importantly, the results confirm that the number of items significantly decrease the precision of temporal reproduction (quantified as the coefficient of variation, CV: mean (item reproduction) divided by standard deviation (item reproduction); F(1, 19) = 45.94, p < 0.001, BF10 > 1000). Conversely, the duration of the sequence did not significantly affect the precision of temporal reproduction (F(1, 19) = 0.51, p = 0.49, BF10 = 0.13). Thus, the number of durations, but not the duration of the sequence itself constitutes the load when it comes to storing multiple durations in working memory.

### Decoding results

Next, we applied common spatial pattern decoding (CSP, Grosse-Wentrup & Buss, 2008; Koles, 1991) on the MEG data to test whether oscillatory dynamics during the stimulus-free retention interval contained information about the number of items and/ or the duration of the sequence. Decoding was applied to induced oscillatory power (after subtraction of the evoked response), in time-frequency bins (2 – 25 Hz, 0.5 – 4 s). Significant decoding performance at the group level was observed only for the number of items (**Error! Reference source not found**.A, left), but not for the duration of the sequence (**Error! Reference source not found**.B). Standard error of the mean for the time x frequency matrices is displayed in **Supplementary Figure 2**. A cluster permutation test revealed a statistically significant cluster spanning all time-frequency bins (**Error! Reference source not found**.A, left). The peak frequency distributions (histograms in **Error! Reference source not found**.A, middle) were of bimodal shape, with one peak in the alpha and one in the beta band. The source projection of the decoding patterns for the whole frequency and time ranges showed sources in occipital, parietal, and motor areas (**Error! Reference source not found**.C). To investigate the cortical dynamics underlying the significant decoding performance, we separated the decoding patterns in the alpha and beta bands before projecting the weights in source space (**Error! Reference source not found**.E). This revealed parieto-occipital sources in the alpha band, versus a more pronounced spread into sensorimotor areas for the beta band. Both were relatively lateralized to the left hemisphere. Lastly, we reconstructed the sources directly from the oscillatory power, rather than from the decoding patterns, which allowed assessing the directionality of the effect: for the alpha band, we observed higher power for 3-item versus 1-item sequences, with a bilateral occipital distribution. In the beta band, power was higher for 1-item compared to 3-item sequences in the motor regions, and higher for 3-items in occipital regions (**Error! Reference source not found**.F), both bilateral. While the effect of beta power in the occipital regions likely reflects harmonics of alpha, the opposite directionality and the different topographical distributions of the modulation of beta power in the sensorimotor areas suggests that it reflects an independent effect. These findings thus argue for the separability of alpha and beta dynamics during working memory maintenance.

### Alpha power mediates the effect of n-items on precision

We addressed the relationship between the precision of reproduction (quantified by the CV, computed for single trials through a resampling and bootstrapping approach, see Methods) and oscillatory power in the alpha and beta bands, obtained from six source labels covering the significant decoding patterns: lateral occipital cortex, inferior parietal cortex, superior parietal cortex, supramarginal gyrus, precentral and postcentral gyri (all bilateral). Alpha power in the bilateral superior parietal cortices, supramarginal gyri, and postcentral cortex showed a significant relationship with CV in the 1-item trials (correlations ranging from −0.057 – −0.068, all p < 0.003; **Figure 2D**, see also **Supplementary Table 1**, **Supplementary Figure 3**): CV decreased with higher alpha power, meaning precision increased. A marginally significant correlation was observed in the inferior parietal areas (r = −0.053, p = 0.006). No significant effects were found in the 3-item trials, nor for the beta band (all p > 0.02). We then tested for a direct relationship between the behavioral precision effects and the differences in oscillatory dynamics for 3-vs. 1-items through mediation (Shrout & Bolger, 2002). Alpha power in bilateral supramarginal gyrus mediated the effect of n-items on precision: (1) The independent variable n-items had a significant effect on the mediator, power (***a*** effect; −0.06 [-0.07 – −0.06], bootstrapped 95% confidence interval). (2) n-items significantly affected the CV (***b*** effect, −0.1 [-0.17 – −0.04]). (3) The indirect path of n-items on CV via power was significant (***a×b*** effect,0.01,]0.0 – 0.01]).

**Figure 2.**
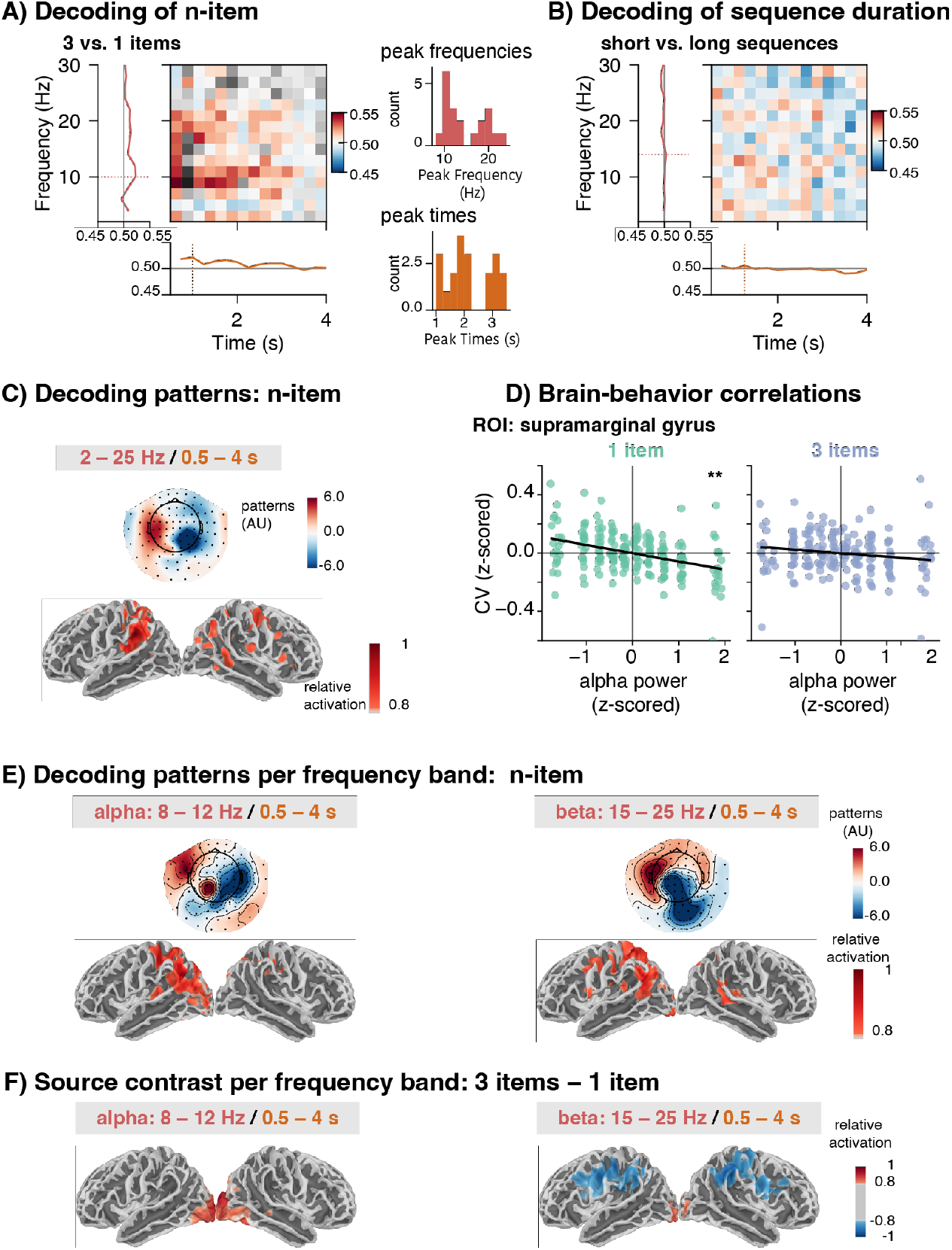
MEG decoding. (Caption for Panels A – B on previous page) C) Sensor topographies and source projections of decoding patterns, averaged across participants. AU = absolute units. D) Brain-behavior correlations. We tested whether the power modulations were directly related to the precision of temporal reproductions per trial. A significant relationship between precision (CV, computed through resampling) and alpha power was found, here depicted for the supramarginal gyrus (sig. only for 1-item; measures were binned only for visualization). E) Sensor topographies and source projections of decoding patterns, separately for the alpha (8 – 12 Hz) and beta (15 – 25 Hz) bands. Decoding patterns in the alpha and beta band overlapped, but were relatively more shifted to occipital/parietal areas for alpha and central areas for beta, and left-lateralized. F) Source contrasts for 3-versus 1-item sequences: power differences projected in source space and thresholded at 80%. While panels A and B show the decoding patterns (red = decoding better than chance), which do not allow to infer the directionality of the effect, the contrast of relative power between 3-and 1-item sequences allows to appreciate the different directions of the effects in the alpha (red = more power for 3 items) and beta bands (blue = more power for 1 item). MEG decoding. A) N-item decoding in sensor space. The number of items was significantly decoded from induced power in the alpha and beta bands during the retention interval using common spatial patterns (CSP). Left: decoding performance during the retention interval, computed in time-frequency bins. The coloured pixels reflect significant decoding performance (decoding accuracy > 0.5; cluster permutation test), and the grey ones non-significant decoding performance. The marginal means reflect average decoding performance across frequency (left, red), and time points (below, orange). Middle: The red and orange histograms show the frequencies and latencies with the highest decoding performance for each participant. B) Decoding of sequence duration (short vs. long). Time-frequency matrix of decoding performance with marginal means reflecting average decoding performance across frequency (left, red), and timepoints (below, orange). No significant decoding performance was observed for sequence duration.

In sum, alpha power in the supramarginal gyrus mediated the effect of the number of items on the precision of temporal reproduction, suggesting that increased working memory load with the number of durations, not with the duration of the sequence.

## Discussion

Sustained representations of durations are required for tasks like duration comparison or reproduction (Van Wassenhove, 2009), but little is known about duration storage in memory. Here, we investigated whether working memory for duration follows similar principles as working memory for sensory items (Bays et al., 2009, 2024; Herbst et al., 2025; Joseph et al., 2016; Ma et al., 2014; Manohar & Husain, 2016). Specifically, we assessed established behavioral and neural markers of working memory maintenance from the sensory domain, namely recall precision and neural oscillatory power during retention, to test whether working memory load for durations is driven by the *number* of durations (n-items) or their duration (i.e. temporal extent) itself. Manipulating n-items and sequence duration orthogonally in a previously published n-item delayed reproduction task (Herbst et al., 2025), we replicated our previous behavioral findings that recall precision decreased with the number of items, but not with sequence duration (**Figure 1**).

During working memory maintenance, a decoding approach applied to MEG data revealed that induced oscillatory dynamics (parieto-occipital alpha and sensorimotor beta) reflected the number of durations (3-vs. 1-item), but not the duration of the sequence (**Error! Reference source not found**.). Furthermore, alpha power statistically mediated the effect of n-items on behavioral precision. A key contribution of this work is the demonstration that time intervals are represented as discrete items in working memory, which extends the principles of sensory working memory to abstract temporal information (Bueti & Walsh, 2009; Gallistel & Gelman, 2000; Herbst et al., 2025; Walsh, 2003).

### Alpha power indexes working memory load

Maintaining three versus one duration in working memory increased alpha power over parieto-occipital areas. Previous studies have established that maintaining multiple distinct visual or auditory items (e.g. letters, faces, or spoken syllables) in working memory is associated with an increase in alpha power measured over parieto-occipital regions, proportional to working memory load (Jensen et al., 2002; Kaufman et al., 1992; Klimesch et al., 1999; Obleser et al., 2012; Scheeringa et al., 2009; Tuladhar et al., 2007). Some studies also reported the opposite direction, but specifically as a relatively short-lived response to visuo-spatial displays (Diaz et al., 2021; Fukuda et al., 2015; Fukuda & Woodman, 2017). The increase of alpha activity is typically interpreted as the disengagement of the dorsal visual stream, to shield the currently retained information from interference by distracting inputs (Bonnefond & Jensen, 2012; X. Chen et al., 2023; Cooper et al., 2003; Jensen & Mazaheri, 2010; Klimesch et al., 2007; Zhou et al., 2023). Our observation extends the signature of alpha power to the maintenance of duration information in working memory. The parietal sources are in line with a previous study reporting load-depended activity modulations in inferior parietal cortex (Teki & Griffiths, 2016), and, more generally, the representation of mental magnitudes in parietal cortices (Bueti & Walsh, 2009), and the representation of durations in chronotopic maps in parietal and motor areas (Harvey et al., 2020; Protopapa et al., 2019).

It has previously been suggested that alpha power could be a signature of spatial attention allocated to more locations with a larger number of items (Foster et al., 2017; Jones et al., 2024). Participants did not mention spatialization as a particular strategy during debriefing, and the most common strategy in timing task is counting (Rattat & Droit-Volet, 2012). Had participants spatialized individual items during retention, subjective distances should increase with item duration according to the classical tau effect (Helson, 1930), which in turn should have surfaced as an effect of sequence duration (similarly for counting), which was not observed.

Critically, alpha power in the supramarginal gyrus statistically mediated the effect of the number of items on recall precision, providing a direct link between the behavioral precision effect and alpha power as an index for working memory load. Furthermore, the absence of a similar effect on alpha power with longer sequence duration, and the inability to decode sequence duration during retention, corroborate the behavioral result that working memory precision depends solely on the number of duration items.

### The role of beta power during working memory retention

Beta power in sensorimotor cortices during retention *decreased* with the number of durations maintained in working memory, which suggests a clear functional distinction from the neural dynamics observed in alpha power over parieto-occipital areas. Beta oscillations are a known neural correlate of working memory, (Bastos et al., 2018; Lundqvist et al., 2016; Miller et al., 2018; Spitzer & Haegens, 2017), with several putative functional roles: top-down inhibition (Bastos et al., 2018; Liljefors et al., 2024), with commonalities and overlaps to alpha dynamics, but also more specifically the maintenance of sequence order (Liljefors et al., 2024; Miller et al., 2018; Salazar et al., 2012). Furthermore, beta power modulations have been reported as neural signatures of interval timing (Kononowicz, 2015; Kononowicz et al., 2018; Kulashekhar et al., 2016) and timing precision (modulated by alpha phase, Grabot et al., 2019; Kononowicz et al., 2020), as well as working memory for duration of a single tactile stimulus (Spitzer et al., 2014).

In line with the inhibitory role of beta, our observation of higher beta power for 1-items here might indicate stronger movement inhibition (Fujioka et al., 2012; Jha et al., 2015; Khanna & Carmena, 2017; Zhang et al., 2008), or enhanced preparation for target encoding (Liljefors et al., 2024), as the maintenance of single items binds less resources than for multi-item sequences. Furthermore, increased complexity in the motor demands (as in the reproduction of several items) could decrease beta power contralateral to the effector side in a motor imagery task (Brinkman et al., 2014), in line with the left lateralization observed in the source reconstructed decoding patterns (Fig. 2B).

With respect to the specific relevance of beta dynamics to timing, our findings point towards a sustained representation of duration in sensorimotor areas used for working memory maintenance. Seminal investigations revealed that working memory maintenance relies on persistent firing of distributed neural populations, both, in areas that encode the respective stimuli (i.e. visual or motor cortices), but also parietal and frontal areas (Bays, 2015; Goldman-Rakic, 1995). Critically, the amplitude of the activity specifically related to the memorized items decreases with load (Sprague et al., 2014), as does recall precision. This decrease can be linked to a more distributed representation of multiple items, suggested to be mediated by beta power (Lundqvist et al., 2016, 2024). Here, the successful decoding of the number of durations in working memory from beta band activity points towards a distributed representation of item durations mediated by beta dynamics.

#### Conclusions

Here, we addressed how sequences of durations are stored in working memory, through behavioral and neural indices of working memory load: recall precision and neural oscillatory dynamics. The number of duration items but not the duration of the sequence had an effect on recall precision, and was decodable from oscillatory power in the alpha and beta bands, pointing towards an abstract representation of ‘duration items’ in working memory. Alpha power increased with load, bridging between durations and other kinds of items. Beta power dynamics differed from alpha, suggesting a specialized neural code for duration.

## STAR METHODS

### EXPERIMENTAL MODEL AND STUDY PARTICIPANT DETAILS

Twenty-three participants (14 women, mean age = 26.2 years, SD = 5.2 years) were recruited at Neurospin. The sample size was chosen in accordance to previous behavioral experiments (Herbst et al., 2025). All participants were right-handed, and had normal hearing and vision, and no self-reported history of audiological or neurological disorders. Participants were naive as to the purpose of the study and received monetary compensation for their participation. Prior to the study, all participants signed a written informed consent in accordance with the Ethics Committee on Human Research at Neurospin: CPP n° 100049 (Gif-sur-Yvette, France), and the Declaration of Helsinki (2013). Three participants were excluded, one because of a problem with the triggers, one had incomplete data, and for the third, no anatomical MRI scan was available. This left 20 participants for the analyses.

Ancestry, race, ethnicity, and socioeconomic status of participants were not recorded, as this information is considered sensitive data under French law and was not covered by the current ethics approval. While the sample characteristics are standard for resource-intensive neuroimaging studies aiming for relatively small, homogeneous samples, they have been criticized for not reflecting the worldwide population (Henrich et al., 2010). Given the targeted age group, participants were likely either university students or young professionals, which may have biased the sample’s socioeconomic status. While there is a priori no reason to assume that basic neural and behavioural markers like the ones assessed here should be affected by these individual characteristics, future research should extend these findings to more diverse populations, particularly a wider age group.

## METHOD DETAILS

### Data and Code Availability

The anonymized MEG data including behavioral markers (BIDS format) are available in a private repository on OpenNeuro (for review only). Upon acceptance, the repository will be made public with a link included in the final version of the paper. The custom code in python is available in a private repository on the OSF, which will be made public upon publication. The study was not formally pre-registered.

### General Procedure and Task

Participants performed an *n-item delayed temporal reproduction* task (Herbst et al., 2025). They were presented with a sequence of one or three “empty” intervals (Fig. 1A), delimited by short pure tones (*encoding*; for details about the stimuli see below). Empty time intervals were used to prevent a maintenance strategy based on auditory features, as opposed to duration. They had to maintain the sequence in memory (*retention*), and, upon a prompt, reproduce the whole sequence by pressing a button for each tone (*reproduction*). Participants received feedback after each reproduction of a sequence.

Before the main experiment, the researcher explained the task to the participants, and provided them with written instructions. Participants then started a training block composed of eight trials. The training was repeated if a participant failed on more than half of the trials (average relative temporal reproduction error over all items in the sequence below −0.5 or above +0.5). Data from the training session were not included in the analyses. The actual experiment consisted of 8 blocks of 36 trials each; each block lasted about 10 minutes. After each block, the MEG recording was stopped to save the data, resulting in a break for the participant. The total MEG session lasted 90 minutes. Each participant was presented with the same conditions, in randomized order as described above. As the design was a repeated-measures within subject design, no blinding was put in place.

### Stimuli

The experimental paradigm was adapted from Experiment 2 in Herbst et al. (2025). All stimuli were created digitally using Psychtoolbox (Brainard, 1997) under MATLAB (2017a, The Mathworks), at a sampling rate of 44.1 kHz. Stimuli consisted of a sequence of pure tones (1 kHz, 50 ms duration including 10 ms onset and offset ramping to avoid onset artifacts), which demarcated the time intervals (hereafter referred to as “items”).

Three sequence durations were tested (1.6 s, 2.4 s and 3.6 s) and the number of duration items was one or three for each sequence. Two-item sequences as used in the previous behavioral study were removed to obtain more trials per condition in the MEG experiment. The order of items within each n-item sequence was randomized. We orthogonalized n-items and the duration of the sequences, resulting in a fixed sequence duration regardless of n-items, but shorter individual items in sequences with more items (Figure 1B). Each block contained all possible sequence types in random order: for the 3-item sequences, six permutations of the order exist for each of the three sequence durations (18 trials), and the three different 1-item sequences were each repeated six times throughout the block, to balance the number of 1- and 3-item sequences.

During encoding, a red fixation dot was displayed on the center of the screen. 0.5 s after the last tone in the sequence, an orange fixation dot signaled the onset of the retention period. The retention period lasted for 4.2 s. This relatively long interval was chosen with the aim to assess the neural dynamics of working memory with MEG during a period free from sensory stimuli. Following the retention period, a green fixation dot prompted participants to reproduce the full sequence. To respond, participants pressed one button on a Fiber Optic Response Pad (FORP, Science Plus Group, DE) with the index finger of their right hand. Participants had unlimited time to initiate their reproduction, and no sounds were played when they pressed the button.

To reproduce the sequence, participants pressed the button as many times as there were tones in the sequence (*e*.*g*. 4 key presses for 3 items demarcated by 4 tones), trying to reproduce each duration as precisely as possible. If the correct number of presses was registered, visual feedback was displayed on the screen 0.5 s after the last button press. Visual feedback was displayed for 1 s, and followed by a variable inter-trial interval sampled from a uniform distribution ranging from 2 s to 3 s. Feedback was given in the form of a visual bar plot, which depicted the relative signed reproduction error for each item in the sequence: (item reproduction – item duration) / item duration. Thus, bars of zero length indicated perfectly accurate reproduction, while bars above and below zero reflected over- and under-reproduction, respectively. Trials in which the participant pressed too early (during retention), or an incorrect number of times were labelled as error trials and the participant was prompted with ‘too early’ / ‘wrong number of clicks’ instead of the feedback described above. These trials were removed from the analyses.

### MEG recording

Prior to the arrival of the participant, an empty room recording was performed for one minute to assess the noise level of the MEG sensors. Before undergoing the MEG recording, participants were equipped with external electrodes, positioned to record the electro-occulogram (EOG, horizontal and vertical eye movements) and -cardiogram (ECG). The positions of the EEG electrodes, four head-position indicator coils, and three fiducial points (nasion, left and right pre-auricular areas) were digitized using a 3D digitizer (Polhemus, US/Canada) for subsequent co-registration with the individual’s anatomical MRI. The MEG recordings took place in a magnetically shielded chamber, where the participant was seated in an armchair under the MEG helmet. The electromagnetic brain activity was recorded using a whole-head Elekta Neuromag Vector View 306 MEG system (Neuromag Elekta LTD, Helsinki) with 102 triple-sensors elements (two orthogonal planar gradiometers, and one magnetometer per sensor location). Participants were instructed to fixate their gaze on a screen positioned in front of them, at about one meter distance. The chamber was dimly lit. Their head position was measured before each recording run (8 in total) using the head-position indicator coils. MEG recordings were sampled online at 1 kHz, high-pass filtered at 0.03 Hz, and low-pass filtered at 330 Hz. A two-minute-long resting state recording (eyes open) was performed after the task, used to compute the noise covariance matrix for source reconstruction.

### Anatomical MRI recordings

To improve the spatial resolution of the source reconstruction, individual high-resolution structural Magnetic Resonance Imaging (MRI) recordings were used. These were recorded on another day, using a Siemens 3 T Magnetom Prisma Fit MRI scanner. Parameters of the sequence were: slice thickness: 1 mm, repetition time TR = 2300 ms, echo time TE = 2.98 ms, and flip angle = 9 degrees.

## QUANTIFICATION AND STATISTICAL ANALYSIS

### Software

The analyses of the behavioral data were conducted using R (version 4.3.3 R Core Team, 2024). The MEG data were analyzed using MNE Python (version 1.8.0, Gramfort et al., 2013, 2014; Larson et al., 2024), transformed to the standardized brain imaging data structure (BIDS, Niso et al., 2018) using MNE BIDS (version 1.6.0, Appelhoff et al., 2019), and analyzed with the MNE BIDS pipeline (version 1.8, https://mne.tools/mne-bids-pipeline/), plus additional custom-written code. FreeSurfer was used for the reconstruction of the MRI surfaces (Dale et al., 1999).

### Behavioral data

The reproduced duration per item was measured as the time between the onsets of the two key presses. We removed reproduction outliers defined as reproductions that exceeded by 3 standard deviations the participant’s mean reproduction for that duration. Recall precision is commonly used in working memory experiments as a continuous measure of working memory load (Ma et al., 2014). To ensure that our measure of precision was not confounded by established biases in reproducing durations with varying magnitudes, we employed the **coefficient of variation (CV)**, a widely accepted index of precision in timing research (Gibbon et al., 1984). We computed CV for bins of items of the same duration, and from the same n-item sequence for each participant as follows:

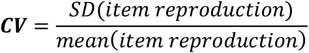

To be in accordance with previous studies, we also computed the **relative reproduction** per item (**relRP**), in the same bins as CV, defined as follows:

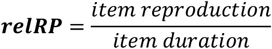

Parallel statistical analyses were performed on **relRP** and **CV**, testing for effects of the number of items (n-items), sequence duration, and their interaction. Random intercepts and slopes were included for all predictors. To assess statistical significance, we computed linear mixed effect models using R’s *lme4* package (Bates et al., 2015), with the Satterthwaite approximation of degrees of freedom, implemented in the *lmerTest* package (Kuznetsova et al., 2016). Additionally, to obtain effect sizes and assess null-effects, we computed Bayes factors (BF) for each predictor in the model using the *bayestestR* package (Makowski et al., 2019) via the Bayesian Information Criterion approximation (Wagenmakers, 2007). The alpha level for assuming statistical significance was set to p < 0.05. In addition, we use Bayes factors to support an unequivocal interpretation of the results. BF represent the relative posterior probability of observing the data under the alternative (i.e., full) and null (i.e., reduced) model, which is informative about whether the inclusion of a given predictor into the model improves the explained variance. For BF we chose the criterion of > 3 and < 0.3 when comparing to models (Lee & Wagenmakers, 2014; Rouder et al., 2009). BF between 0.3 and 3 reflect inconclusive outcomes. In Table 1, we report F- and p-values, as well as BF.

### MEG preprocessing

To remove artifacts from the recorded data, we used a standardized preprocessing routine, implemented in the MNE-BIDS-pipeline, with the following steps: noisy or flat sensors were identified visually and marked for exclusion. Next, environmental artifacts were removed from the raw data, using signal source separation (SSS or Maxwell-filter, Taulu & Kajola, 2005), which also interpolates the bad sensors. Head position coordinates were read from the first of the eight recording runs, and all other runs were spatially aligned to these coordinates. The data were then filtered with a low-pass filter of 160 Hz, and a notch-filter at 50, 100, and 150 Hz to remove line noise. No additional high-pass filter was applied (0.03 Hz used during recording). Furthermore, we resampled the data to a sampling frequency of 250 Hz to speed up the subsequent computations.

Next, the data were cut into epochs of −5 s to 10 s around the onset of the retention interval, and epochs with peak-to-peak amplitudes exceeding 50000 fT in the magnetometers were rejected. We deliberately used no baseline for the time-domain data, as the interval preceding the retention interval contains the evoked activity from the last tone presented during the encoding phase, and the encoding phase itself varies in duration because of the different sequence durations. Independent component analysis (ICA) was performed on the remaining epochs, with an additional 1 Hz high-pass filter applied only for that purpose. For the detection of artefacts related to the electro-occulogram (EOG) and -cardiogram (ECG), we used an inbuilt routine in MNE python, which finds a participant’s typical EOG and ECG activity recorded with the external electrodes, and returns the ICA components that correlate with these typical events. Thresholds for returning the ICA components were set to: 0.1 for the ECG (cross-trial phase statistics, Dammers et al., 2008), and 4.0 for the EOG (z-score).

After ICA cleaning, we rejected epochs with peak-to-peak amplitudes exceeding 10,000 fT (magnetometers) and 20,000 fT/cm (gradiometers). In the following, we report only the analyses of the 102 magnetometers, to reduce the dimensionality of the data and simplify the interpretation of the results and topographies. On average, we obtained 271.7 clean epochs per participant (SD = 13.45, minimum: 246, maximum: 288).

### Anatomical MRI preprocessing and forward model computation

FreeSurfer’s “recon-all” function was used to reconstruct the cortical surfaces of each participant’s brain. Individual surfaces were mapped to FreesSurfer’s ‘fsaverage’ template brain, for later averaging. Single-layer individual head models were computed using the boundary element method (watershed BEM), from each participant’s aMRI. The coregistration of the MEG data with the individual’s structural MRI was carried out by manually realigning the digitized fiducial points with the anatomical landmarks of the head surface reconstructed from the MRI (using MNE Python’s command line tools). Second, an iterative refinement procedure was used to realign all digitized points with the individuals’ scalp. We then computed a surface-based source space with 4098 candidate dipoles per hemisphere, spaced approximately 4.9 mm apart (24 mm^2^ per source area; recursively subdivided octahedrons ‘oct6’), a minimal distance of 5 mm from the inner skull. The final forward models were derived from this source space, the co-registration matrices, and the boundary element models.

### Common spatial pattern decoding

To investigate how duration sequences are represented in working memory and identify the factors contributing to memory load, we focused our MEG data analysis on the retention interval. During this period devoid of sensory inputs, we examined how sequences varying in length and duration modulate *induced oscillatory activity* (Tallon-Baudry et al., 1996), in line with established neural dynamics of working memory (Miller et al., 2018). We employed a decoding approach using Common Spatial Patterns (CSP) in combination with logistic regression, implemented as part of the MNE BIDS pipeline. CSP is recommended for the analysis of induced oscillatory activity (Grosse-Wentrup & Buss, 2008; Koles, 1991). It identifies spatial filters that maximize the variance of one class of signals (e.g., sequences with a three versus one item), while minimizing the variance of other classes. This allowed us to effectively classify brain activity patterns associated with varying memory load.

We performed the CSP spatial filtering in linearly spaced time-frequency bins, ranging from 2 – 30 Hz (bin width of 2 Hz, 14 bins) and 0.5 – 4 s following retention onset (bin width of 0.25 s, 14 bins). Specifically, the filtering and decoding was run on each trials’ covariance matrix computed on single trial data, band-pass filtered in the range of the desired frequency bin, after subtracting the evoked response to focus on induced activity. Since no time-frequency transformation is performed here, there was also no additional baseline-correction. For decoding, we used logistic regression (solver: “liblinear”) with a 5-fold cross validation (shuffled splits), and quantified decoding performance as the area under the receiver-operator curve averaged across the folds.

To assess in which frequency bands and at which time points we could significantly decode the number of items or the sequence duration at the group level, we performed cluster-permutation tests on the decoding accuracy values across time-frequency bins, using an initial cluster forming threshold of p < 0.01, a final cluster selection threshold of p < 0.05, and 5000 permutations.

### Source reconstruction

To further investigate which brain areas contribute to the significant decoding accuracy, we reconstructed the sources of the decoding patterns (Westner & King, 2023)^1^. Here, we used the decoding weights (recomputed now for all epochs jointly without cross-validation), and transformed them to the corresponding patterns across sensors (group average depicted in Figure 2). Topographical patterns are closer to an interpretation as neurophysiological activity compared to the classifier weights (Haufe et al., 2014). We then computed linearly constrained minimum variance spatial filters (lcmv-beamformers, Van Veen et al., 1997) from the epoched and band-pass filtered data, and a noise covariance matrix from the resting state recording, and projected the decoding patterns (back-transformed to covariance matrices) to the source level. Finally, we morphed the individual source activity to a template brain for averaging (fsaverage). We performed the source reconstruction for the complete range of frequencies identified as statistically significant, and second, split into the canonical alpha (8 – 12 Hz) and beta (15 – 25 Hz) frequency bands. Finally, to investigate the directions of the effects, we also projected the frequency-band power difference (covariance matrix computed from band-pass filtered epochs) for 1-item and 3-item sequences to source space using minimum-norm estimates (dSPM, Dale et al., 2000).

### Brain-behavior relationships

To further assess whether the neural dynamics identified as correlates of working memory load for durations directly relate to behavioral reproduction precision, we extracted single trial power in the alpha (8 – 12 Hz) and beta (15 – 25 Hz) bands from the occipital, parietal, and central regions, in which we found significant decoding (labels: lateraloccipital, inferiorparietal, superiorparietal, supramarginal, precentral, postcentral from the ‘aparc’ atlas (Destrieux et al., 2010)). We created functional labels for each participant per hemisphere, by sub-selecting the 15 % of voxels with the highest activity in each label (across all conditions). Selecting only the most activated voxels balanced the need for robust labels, despite individual differences, with the goal of maximizing the signal-to-noise ratio by excluding inactive voxels. We then computed the average power for the alpha and beta band for each label, by applying the Hilbert transform to bandpass-filtered data from each epoch to obtain complex time series, and squaring the absolute values averaging across voxels and time points (0.5 – 4 s) to obtain oscillatory power.

To obtain precision *for each trial*, a measure which is per se defined *across* trials, we applied a resampling and bootstrapping technique (Tibshirani & Efron, 1993). Per participant and n-items, we randomly selected 20 trials (without replacement), and computed one value of CV and average power for this sample. We repeated this 1000 times. To test for significant correlations between power and CV, we computed correlations between CV and power across these 1000 values, and compared them to a null-distribution, for which we shuffled the indices of the 1000 samples and computed the correlation, repeating this shuffling 1000 times to obtain a null-distribution of 1000 correlation values (**Supplementary Table 1**, **Supplementary Figure 3**). CV was z-scored across all participants and trials for model stability, and power values were averaged for the left and right hemispheres and z-scored per individual, as recommended by e.g. Grandchamp & Delorme (2011). In order to compute p-values, we counted how many correlation values obtained from the null distribution were as large or larger than the true correlation value. We set a threshold of p < 0.004 (p < 0.05 with Bonferroni correction for 12 comparisons). We targeted only CV for this analysis (and not relRP), as it provides a bias-free measure of recall precision in particular with respect to wide range of durations used.

### Mediation analyses

We also assessed whether oscillatory power in the retention window mediates the effect of n-items on CV (Baron & Kenny, 1986; Benwell et al., 2018; Memon et al., 2018; Muller et al., 2005; Shrout & Bolger, 2002), i.e. whether the neural dynamics play the hypothesized statistical role to index working memory load caused by the number of items. To test for the presence of a statistically significant mediation, we followed the steps described in Shrout & Bolger (2002). We used the same resampling technique as above to compute power and CV for 1000 samples for each participant (500 per n-item). We computed linear mixed effect models using the *mixedlm* function from the *statsmodels* package for python (Seabold & Perktold, 2010). The models were twofold: (1) regressing power on n-items (***a*** effect) and (2) regressing CV on power (***b*** effect) and n-items (***c’*** effect). Both models had a random intercept and random slopes across participant for each fixed effect. From these models, we estimated regression coefficients for the three effects, and repeated the sampling 5000 times to obtain reliable 95% bootstrap confidence intervals (Tibshirani & Efron, 1993). The mediation was quantified as the product of the ***a × b*** effect. If the confidence interval for this effect does not include zero, a significant mediation can be assumed (Shrout & Bolger, 2002). Results are depicted in **Supplementary Table 2** and **Supplementary Figure 4**.

## Acknowledgments

This project has received funding from Agence Nationale de la Recherche ‘AutoTime’ (ANR-16-CE37-0004-04) granted to Virginie van Wassenhove, Agence Nationale de la Recherche meegBIDS.fr (ANR-19-DATA-0023), granted to Alexandre Gramfort (Coordinator), Sophie Herbst, Maximilien Chaumon, and Virginie van Wassenhove (collaborators), and from the European Union’s Horizon 2020 research and innovation program under grant agreement No. 101017727, FET Experience granted to Virginie van Wassenhove.

NeuroSpin, NeuroPSI, UNICOG, Cognition & Brain Dynamics, and the Mind Lab are part of the DIM C-BRAINS, funded by the Conseil Régional d’Ile-de-France. We would also like to thank the NeuroSpin MEG platform staff, and in particular Leila Azizi, for their support during data acquisition. We thank Yunyun Shen for enriching discussions about the study, both theoretically and empirically.

## Supplementary Materials

**Supplementary Figure 1.**
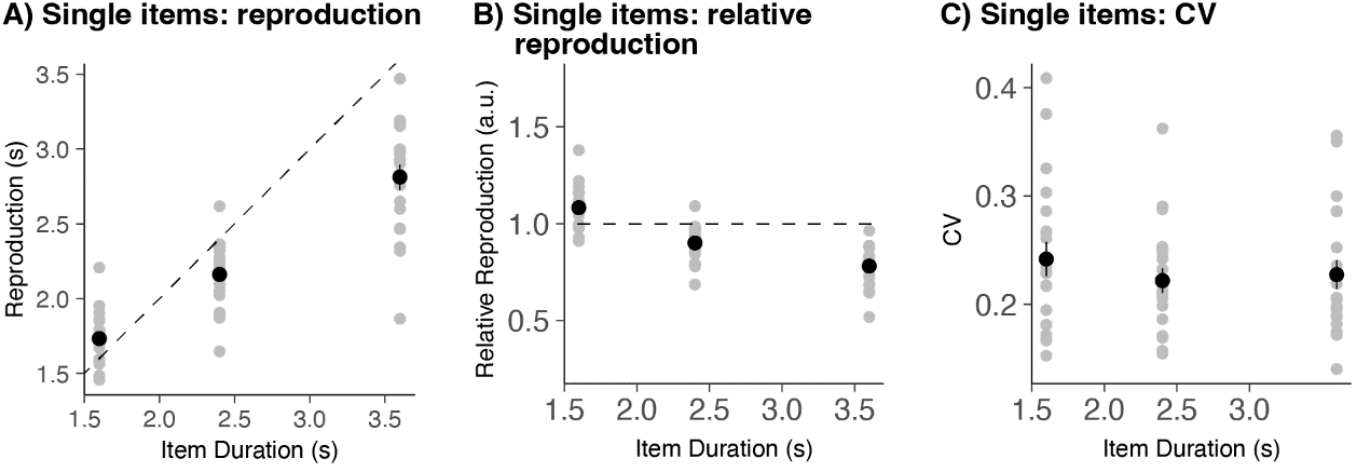
Reproduction of single items. **A:** Reproduction of single items (in seconds), as a function of item duration. Shorter items were over-produced, and longer items under-produced, relative to their objective value. This reflects a regression of the mean effect, typically found in timing studies. **B:** Relative reproduction (reproduced duration divided by item duration), as a function of item duration. **C:** Coefficient of Variation (CV) of single item reproduction. There was no significant difference between the CV of different item durations, indicating that scalar variability was observed in this task.

**Supplementary Figure 2.**
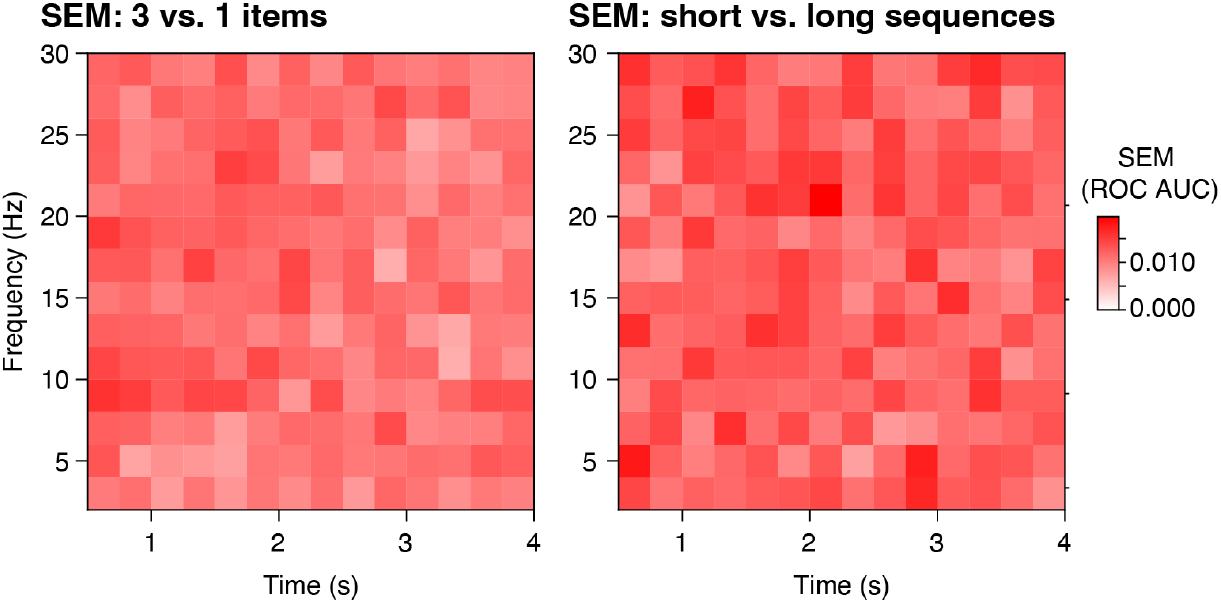
Standard error of the mean (SEM) for the average decoding scores depicted in Figure 2.

**Supplementary Figure 3.**
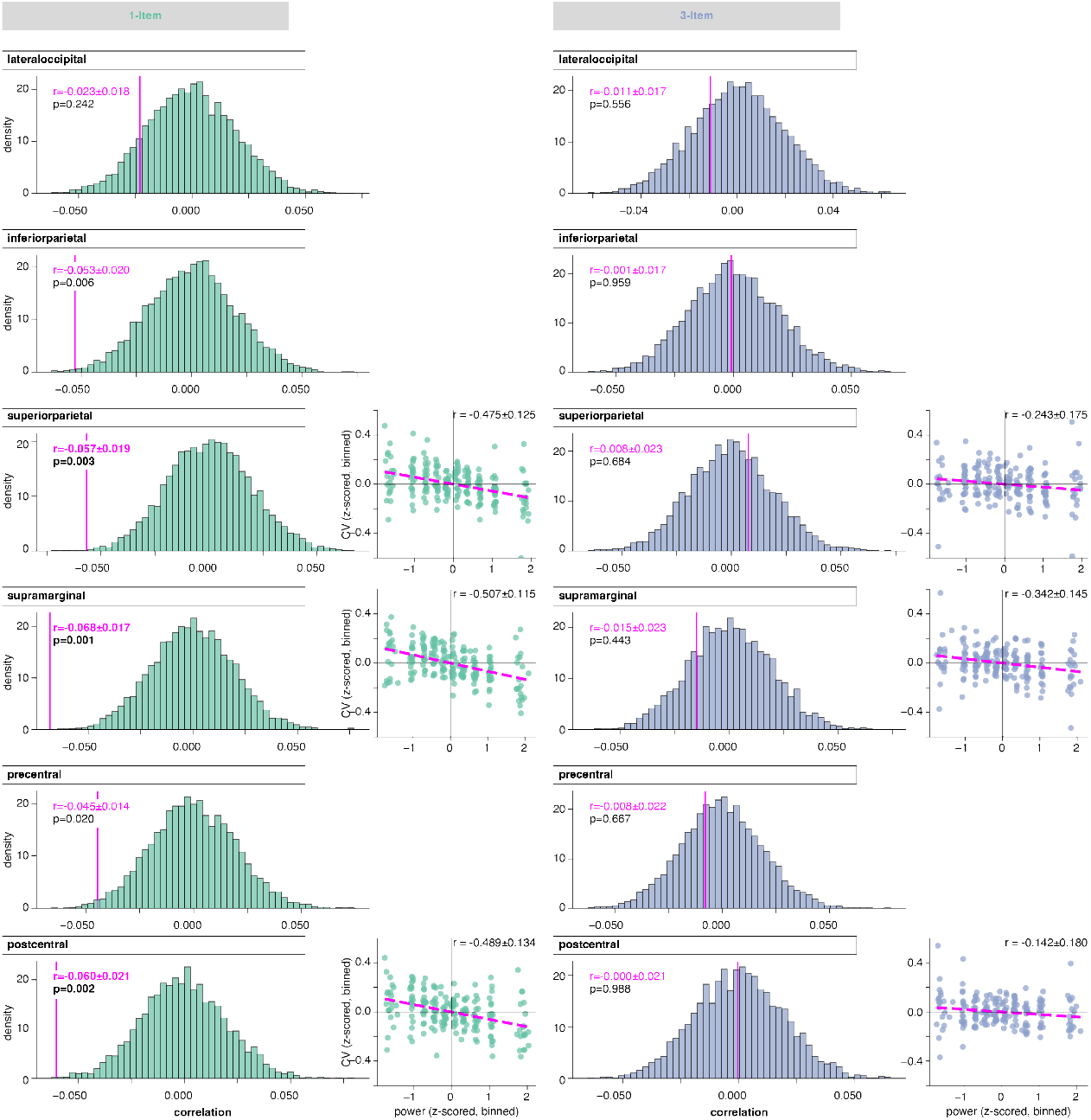
Brain-behaviour correlations between alpha power extracted from source labels and CV. Left: 1-item, Right: 3-item. CVs were obtained through resampling of trials, and then correlated with power values obtained from the same samples. The vertical magenta line depicts the correlation value r ± 95% bootstrap confidence intervals. Green and blue histograms display a null-distribution of correlations calculated after shuffling the indexing between CV and power samples, also used to compute p-values. Dashed magenta squares highlight significant correlations (p < 0.004, Bonferroni correction). Scatter plots show the correlation between power and CV for the significant labels; samples were binned only for display. The r values written in the top right corner of the correlation plots correspond to the binned samples and are therefore higher than the r values obtained from the repeated sampling.

**Supplementary Table 1.** Brain-behaviour correlations between power extracted from source labels and CV, for the alpha (top half) and beta frequency bands (bottom half), separated for n-items. CVs were obtained through resampling of trials, and then correlated with power values obtained from the same samples. Values are correlation value r ± 95% bootstrap confidence intervals. P-values were obtained from a null-distribution of correlations calculated after shuffling the indexes between CV and power samples. Significant correlations are marked in bold font (p < 0.004, Bonferroni correction).

| frequency band | label | 1-item |  | 3-item |  |
| --- | --- | --- | --- | --- | --- |
| ALPHA |  | correlation<br>CV × power<br>± bootstrapped CI | p-value | correlation<br>CV × power<br>± bootstrapped CI | p-value |
|  | lateral occipital | -0.023 ± 0.018 | 0.24 | -0.011 ± 0.017 | 0.56 |
|  | inferior parietal | -0.053 ± 0.020 | 0.006 | -0.001 ± 0.017 | 0.96 |
|  | **superior parietal** | **-0.057 ± 0.019** | **0.003** | 0.008 ± 0.023 | 0.68 |
|  | **supramarginal** | **-0.068 ± 0.017** | **0.001** | -0.015 ± 0.023 | 0.44 |
|  | precentral | -0.045 ± 0.014 | 0.020 | -0.008 ± 0.022 | 0.67 |
|  | **postcentral** | **-0.060 ± 0.021** | **0.002** | 0.000 ± 0.021 | 0.99 |
| BETA | label | 1-item |  | 3-item |  |
|  |  | correlation<br>CV × power<br>± bootstrapped CI | p-value | correlation<br>CV × power<br>± bootstrapped CI | p-value |
|  | lateral occipital | 0.006 ± 0.018 | 0.77 | -0.015 ± 0.017 | 0.43 |
|  | inferior parietal | -0.000 ± 0.022 | 0.99 | -0.028 ± 0.023 | 0.15 |
|  | superior parietal | -0.018 ± 0.022 | 0.34 | 0.004 ± 0.020 | 0.83 |
|  | supramarginal | -0.025 ± 0.023 | 0.19 | -0.033 ± 0.023 | 0.09 |
|  | precentral | -0.022 ± 0.022 | 0.27 | -0.008 ± 0.019 | 0.69 |
|  | postcentral | -0.021 ± 0.023 | 0.28 | -0.001 ± 0.024 | 0.96 |

**Supplementary Table 2.**
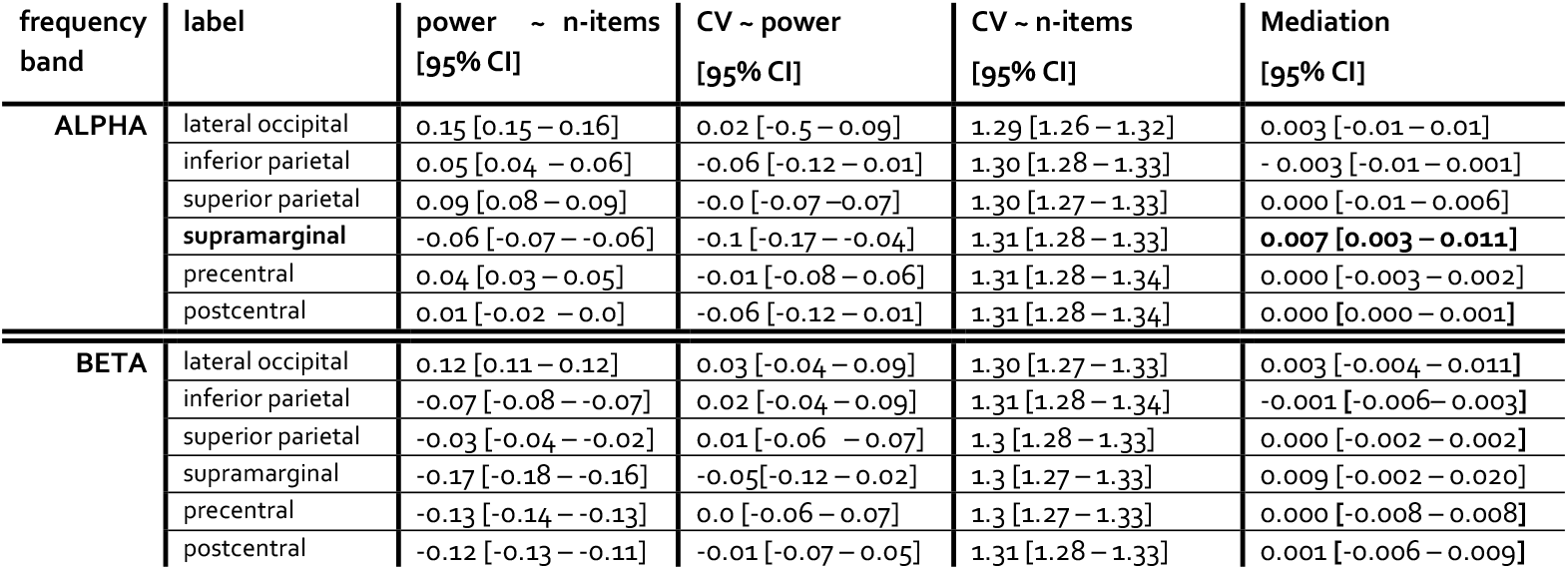
Mediation analyses, conducted for alpha (top half) and beta power (bottom half) in source labels (Shrout & Bolger 2002). CV values were obtained through resampling of trails. Values depict regression weights (linear mixed effect models with fixed effects and random slopes) and 95% bootstrapped confidence intervals. We tested for effects of n-items on power (***a***, 2^nd^ column), power on CV (***b***, 3^rd^ column), n-items on CV (***c’***, 4^th^ column), and the indirect, or mediation effect of n-items on CV via power (5^th^ column). Significant mediation (not corrected for multiple comparisons) was found in the alpha band supramarginal gyrus.

**Supplementary Figure 4.**
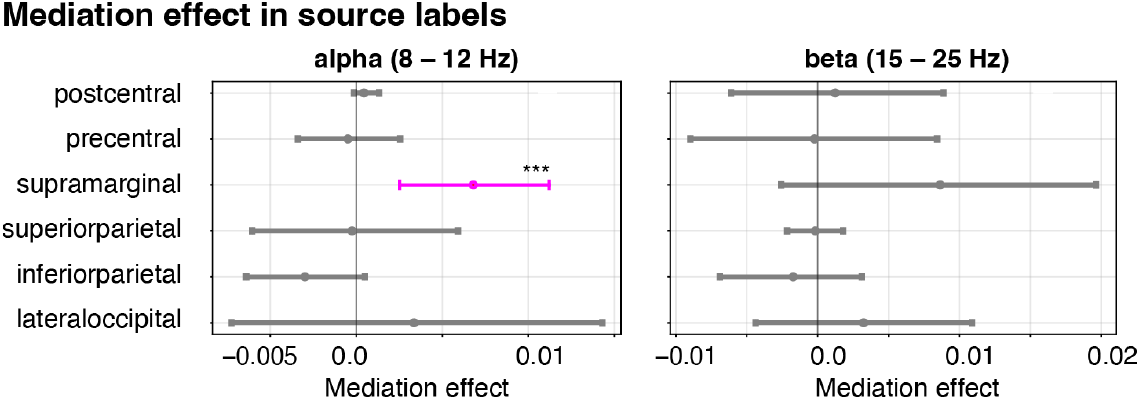
Mediation analyses. Horizontal lines depict the indirect, or mediation effect of n-items on CV via power (5^th^ column in Table S2). Significant mediation (not corrected for multiple comparisons) is highlighted in magenta, and was found in the alpha band in the supramarginal gyrus.

## Footnotes

1 *https://github.com/britta-wstnr/source_decoding*

